# Extensive adaptive immune response of AAVs and Cas proteins in non-human primates

**DOI:** 10.1101/588913

**Authors:** Puhao Xiao, Raoxian Bai, Ting Zhang, Yin Zhou, Zhigang Zhou, Yan Zhuo, Nannan Gong, Jie Liu, Shuaiwei Ren, Ruo Wu, Bin Shen, Shangang Li, Yuyu Niu, Weizhi Ji, Yongchang Chen

## Abstract

The CRISPR-mediated Cas system is the most widely used tool in gene editing and gene therapy for its convenience and efficiency. Delivery of the CRISPR system by adeno-associated viruses (AAVs) is currently the most promising approach to gene therapy. However, pre-existing adaptive immune responses against CRISPR nuclease (PAIR-C) and AAVs has been found in human serum, indicating that immune response is a problem that cannot be ignored, especially for *in vivo* gene correction. Non-human primates (NHPs) share many genetic and physiological traits with human, and are considered as the bridge for translational medicine. However, whether NHPs have same PAIR-C status with human is still unknown. Here, macaques (rhesus and cynomolgus), including normal housed and CRISPR-SpCas9 or TALENs edited individuals, were used to detect PAIR-C which covered SaCas9, SpCas9, AsCas12a and LbCas12a. Dogs and mice were also detected to expand the range of species. In addition, pre-existing adaptive antibodies to AAV8 and AAV9 were performed against macaques of different ages. The results showed that adaptive immunity was pre-existing in the macaques regardless of Cas proteins and AAVs. These findings indicate that the pre-existing adaptive immune of AAV-delivered CRISPR construction and correction system should be concerned for *in vivo* experiments.

## Introduction

The existing popular gene editing technologies mainly include zinc finger nuclease (ZFN), transcription activator-like effector nucleases (TALENs), and clustered regular interspaced short palindromic repeats (CRISPR) [1–3]. Through these techniques, target genes can be knocked out or knocked in, and animal models of diseases caused by gene mutations can be obtained[4–7]. Due to its convenient design, simple operation and high efficiency, CRISRP system has become the most used gene editing system[8]. With the further development of the CRISPR system, it has begun to be applied to preclinical trials to treat or repair hereditary diseases caused by gene mutations[9]. Currently, there are two main methods for applying the CRISPR system to therapeutic preclinical experiments. One is *in vitro* editing. For example, stem cells are edited *in vitro*, and the repaired cells are returned to diseased tissues and organs[10–12]. The other is *in vivo* editing, using vectors to deliver the CRISPR system to the diseased area by intravenous, intramuscular or intraperitoneal injection[13–15]. However, due to the inherent nature of the CRISPR system, its safety such as off-target[16–18], immune response[19–21] and toxicity[22–24] have not been well resolved, making it a huge challenge for clinic application.

The CRISPR system is mainly derived from bacteria and archaea [25], which often invade organisms as pathogens, and are recognized as an alien by the immune system of these organisms, activates lymphocytes in the body to produce corresponding effector cells, and removes them[26]. It is difficult for humans or animals to avoid contact with microorganisms under normal living conditions, and antibodies specific to these pathogens are usually present in the body. Several research groups have found the pre-existing adaptive immune responses against CRISPRCas9 nuclease (PAIR-C) are extensively existing in human, which focus on specific humoral and cellular immunity against SaCas9 and SpCas9[19, 20]. These findings raise concerns on the safety of the *in vivo* CRISPR editing system. Therefore, systematic preclinical assessment of PAIR-C using proper laboratory animals is necessary.

AAVs have been widely used as a vector for gene therapy in preclinical and clinical trials because it is not pathogenic. However, human and Non-human primates (NHPs), as natural hosts for AAVs, have been generally found to neutralize antibodies (NABs) against AAVs, which affect transduction efficiency *in* vivo[27, 28]. Studies have shown that the capsid protein of AAVs can also be recognized by T cells, and cellular immunity of AAVs cannot be ignored[29, 30].

NHPs, such as macaques (rhesus and cynomolgus), are closely related to human beings, have unique advantages in modeling clinic diseases, as well as preclinical safety and efficacy assessment[31]. The pre-existing adaptive immune responses against Cas proteins and AAVs assay in macaques could be helpful to promote clinical application of gene editing technologies. Here, we detected the pre-existing adaptive immune against AAV8 and AAV9 NABs in the serum of macaques. Meanwhile, our team previously obtained several gene-editing macaque models by injecting TALENs plasmid or CRISPR-Cas9 system in one-cell embryos[4, 7]. These models are quite valuable for evaluate the PAIR-C in gene edited and normal wild-type monkeys. In addition, adaptive antibodies against Cas9 proteins have been found in humans[20], but it is not clear whether other species are the same. Organisms have been found to have adaptive antibodies against type II Cas9 nucleases, while type V Cas12a (Cpf1) nucleases have not been reported[20, 32]. To verify whether there are preexisting antibodies against Cas proteins in macaques and other species, we designed prokaryotic expression vectors to express SaCas9, SpCas9, AsCas12a (AsCpf1) and LbCas12a (LbCpf1) proteins in *Escherichia coli*, and purified the four proteins by Ni-NTA affinity chromatography column. Serum samples of macaques, dogs, and SPF mice were isolated, and the pre-existing adaptive immune was assayed by western blot and ELISA.

## Results

### NABs against AAVs are widespread in macaques

As an efficient and safe carrier, AAVs play an important role in gene therapy. *In vivo* NABs against AAVs determine the efficiency of AAVs transduction. We used enzyme-linked immunosorbent assay (ELISA) to detect antibody levels of AAV8 and AAV9 in macaques (n = 84), and found that AAV8 and AAV9 antibodies were widely present in macaques and correlated with age (Fig 1a and 1b). The results showed that the antibody level in the macaques (1-4 years old) was relatively low, and the antibody level in the middle-aged macaques (5-9 years old) was significantly increased, which was significantly different from that of the one-year-old macaques (p<0.01). The levels of AAV8 and AAV9 antibodies in aged monkeys (10-12 years old) were lower than those in middle-aged monkeys, but still higher than that in young monkeys. The results show that NABs to AAV8 and AAV9 are widely present in macaques of all ages, and we speculate that applying them to gene therapy at the living level may cause a certain degree of immune response.

**Fig 1.**
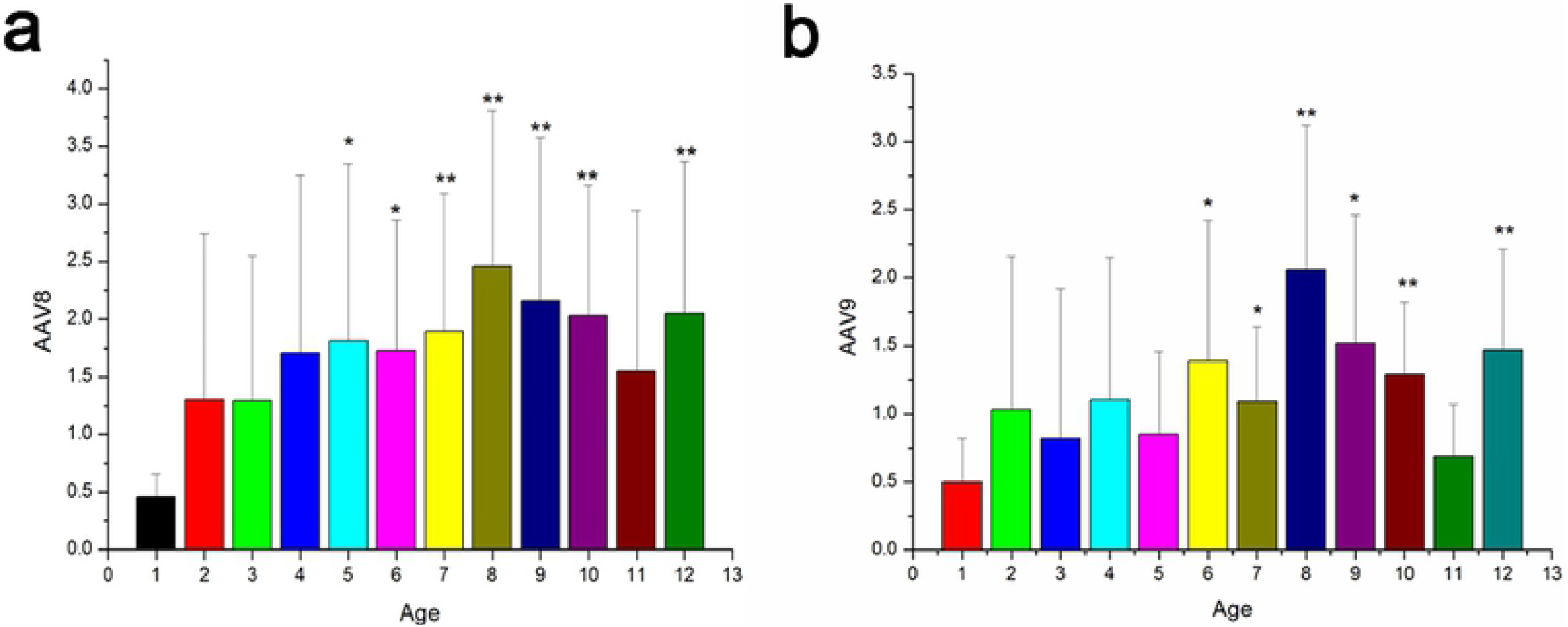
Detection of AAVs antibodies in serum of monkeys. Seven macaques were selected from each age group of 1-12-year-old macaques, and a total of 84 samples were tested by ELISA. (a) is the result of detecting the antibody level of AAV8; (b) is the result of detecting the antibody of AAV9. One-way ANOVA was performed with reference to age group 1 year. * p < 0.05, ** p < 0.01.

### SaCas9, SpCas9, AsCas12a, LbCas12a were synthesized and purified

To detect antibodies against Cas proteins in serum, we selected SaCas9, SpCas9 nucleases in Type II of Class 2 in the CRISPR system, and AsCas12a and LbCas12a nucleases in Type V. In order to obtain SaCas9, SpCas9, AsCas12a, and LbCas12a proteins, we designed primers containing nuclear localization signals (NLS) for the above four protein sequences according to the plasmid published by Zhang Feng laboratory[33–35], and constructed prokaryotic expression vectors pET28b-SaCas9, pET28b-SpCas9, pET28b-AsCas12a and pET28b-LbCas12a (S1a Fig). After double digestion and sequencing verification, the four vectors were transformed into Rosetta (DE3) *E. coli* competent cells, respectively (S1b Fig). The strain carrying the target sequence was induced by IPTG to produce the target protein (S1c Fig). Subsequently, the bacteria were disrupted by sonication and the four proteins of interest were purified by immobilized metal affinity chromatography (IMAC). The purified proteins were detected by polyacrylamide gel electrophoresis (PAGE) (S1d-g Fig).

### The macaque serum PAIR-C is extensively existed regardless of gene editing

In order to detect antibodies against Cas proteins in macaques, we selected 24 healthy wild-type experimental macaques to collect serum and divided into four groups according to age: under 1 year old (n = 3), 1-3 years old (n = 8), 4-8 years old (n = 7) and over 9 years old (n = 6). At the same time, 11 CRISPR-SpCas9 edited macaques (6 positive and 5 negative) and 10 TALENs edited macaques (5 positive and 5 negative) were collected for further analysis (S1 Table). The four proteins of SaCas9, SpCas9, AsCas12a, and LbCas12a were purified and subjected to detect PAIR-C level in above monkeys through western blot. The diluted sera were used as a primary antibody to bind Cas proteins, and anti-monkey IgG H&R was used as a secondary antibody to detect IgG capable of binding to Cas proteins. The antibody was detected by automatic exposure to protein chemiluminescence imager. We found that except for the three macaques under the age of one, most of monkey serum samples contain antibodies to the four Cas proteins (S1 Data, S4 Table). The antibody occurrence rate for SaCas9, SpCas9, AsCas12a, LbCas12a were 97.78%, 95.57%, 95.57%, and 97.78%, respectively (Table 1). These results indicate that the macaque serum antibodies are extensively existed regardless of gene editing, and baby monkey less than one year are relatively mild.

**Table 1.** Frequency of humoral immune response to Cas9 and Cas12a proteins.

|  | **SaCas9** | **SpCas9** | **AsCas12a** | **LBCas12a** |
| --- | --- | --- | --- | --- |
| **Monkey** | **97.78%** | **95.57%** | **95.57%** | **97.78%** |
| **Dog** | **100%** | **100%** | **100%** | **100%** |
| **Mouse** | **92.86%** | **92.86%** | **92.86%** | **92.86%** |

### The serum PAIR-C in dogs and mice is also existed at different levels

To determine and compare the status and levels of PAIR-C in monkeys, we selected 10 dogs (S2 Table) and 16 SPF mice for comparison (S3 Table). Based on the results of western blot, we found that the presence of adaptive antibodies to these four proteins is also prevalent in dog serum as in monkey (Data S1). Unlike previous predictions, antibodies to these four nucleases were also detected in SPF mice (Data S1). Statistical results indicate that all dog serum samples were present for antibodies to these four Cas proteins (Table 1, S5 Table). Moreover, 15 of the 16 SPF mice detected antibodies against these four Cas proteins (Table 1, S6 Table). Due to the exposure duration for monkeys and dogs are quite shorter than for mice, it indicated the real PAIR-C levels in mice are different with dogs and monkeys.

### Quantitative comparison of antibody level

To compare antibody levels in the same species and in different species, we further adopted ELISA to quantify antibody levels in different serum samples. Since the ELISA indirect method requires high-purity antigen (Cas proteins) for coating, it is difficult to ensure the accuracy of the ELISA results by one-step purification of the protein by a Ni-NTA affinity chromatography column[36, 37]. To meet the needs of the experiment, we chose the most commonly used SpCas9 nuclease, which is also the effector of the CRISPR in gene editing monkeys, and purchased the SpCas9 recombinant protein. According to ELISA results, we found that antibody levels in macaques tend to increase with age and remain relatively stable after adulthood (Fig 2a). One-way analysis of variance (ANOVA) of wild-type monkeys, dogs, and mice showed that there were significant differences between monkeys and mice, as well as between dogs and mice. But there was no significant difference between monkeys and dogs (Fig 2b). We also performed a one-way ANOVA of the macaque positive and negative groups edited by CRISPR-Cas9 and TALENs, with no significant differences (Fig 2c and 2d). At the same time, the wild-type population was compared with the monkey population edited by CRISPR-Cas9 and TALENs, and no significant difference was found (Fig 2e).

**Fig 2.**
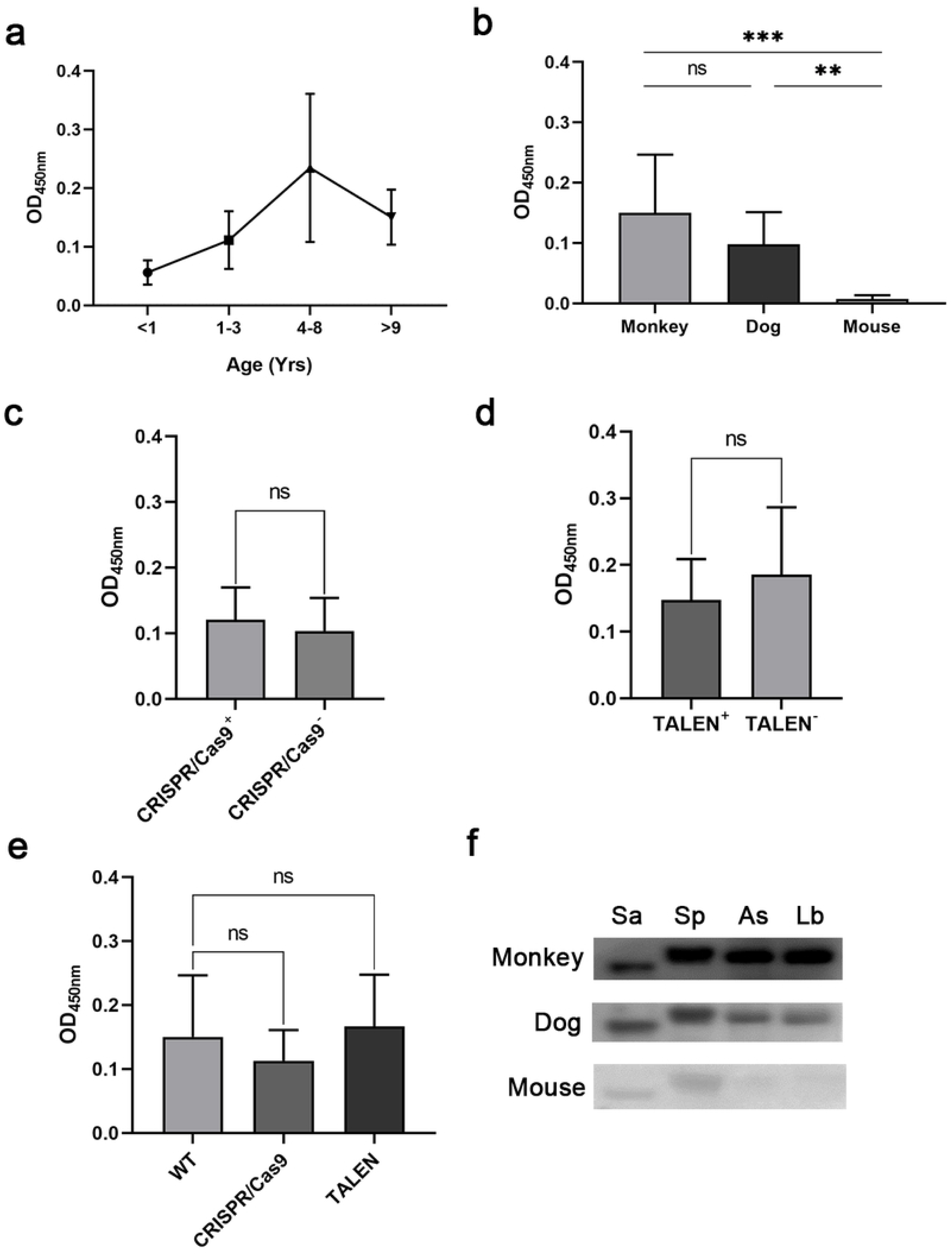
Detection of Cas proteins antibodies in serum of monkeys. (a) Antibody levels of SpCas9 in serum of wild-type macaques of different ages were detected by ELISA at OD450nm. (b) ELISA results for SpCas9 antibodies in wild-type macaque, dog and SPF mouse sera. (c) One-cell embryo stage was injected with the CRISPR/Cas9 system, and the Sanger sequencing test edited (+) and unedited (-) macaque ELISA test results, and there was no significant difference. (d) One-cell embryo stage was injected with TALENs plasmid, and the Sanger sequencing test edited (+) and unedited (-) macaque ELISA test results, and there was no significant difference. (e) Elisa results of wild-type macaques, one-cell embryos injected with CRISPR/Cas9 system, and one-cell embryos injected with TALENs. There was no significant difference in the sera of these three groups of macaques against the SpCas9 antibody. (f) The median Elisa results of macaques, dogs and mice were selected for western blot analysis and the results were simultaneously exposed under protein chemiluminescence imager. **P <0.01, ***P <0.001, ns, not significant, one-way ANOVA with T-test.

In consideration of species difference and sensitive of exposure during western blot, and to verify the results of ELISA and western blot, we ranked the ELISA values of macaques, dogs and mice, and selected the corresponding median serum samples for western blot analysis of four Cas proteins. The results were obtained under the condition of simultaneous exposure (Fig 2f, S2 Fig). Under simultaneous exposure, the results showed that the bands of monkey and dog were significantly stronger than mouse.

## Discussion

AAV belongs to the *Dependoparvovirus* genus, *Parvoviridae* family. Due to the high safety of AAV, it has become the preferred platform for gene delivery[38, 39]. AAVs are present in a variety of spinal animals, including humans and NHPs. Although AAVs itself does not cause disease, as a carrier of gene delivery *in vivo*, inflammatory reactions and immune barriers are inevitable problems. At present, AAVs are mainly used in the treatment of eyes and brain in clinical experiments[39]. AAV8 and AAV9 can target a variety of muscles throughout the body. When using AAV systemic treatment, how to avoid immune system monitoring not only directly affects the effect of treatment, but also determines the difficulty of re-dosing.

The CRISPR/Cas system is an acquired immune system among most bacteria and all archaea, in which SaCas9 is derived from *Staphylococcus aureus*, SpCas9 is derived from *Streptococcus pyogenes*, AsCas12a is derived from *Acidaminococcus sp*., and LbCas12a is derived from *Lachnospiraceae bacterium*[33–35]. Generally, humoral immunity requires antigen stimulation to produce antibodies. These microorganisms are often recognized by the immune system as foreign substances and stimulate B lymphocytes to produce antibodies. In this process, Cas or Cas-like proteins are a potential antigen that activates the immune system to produce antibodies. To verify the safety of the CRISRP system, we detected the presence of adaptive antibodies against SaCas9, SpCas9, AsCas12a and LbCas12a in macaques, dogs and SPF mice serum by western blot. To further quantify the level of antibodies, we again detected antibodies against SpCas9 protein in the macaques, dogs and mice serum by ELISA. These results indicate that pre-existing adaptive antibodies to these four proteins are ubiquitous in different species, but antibody levels in SPF mice are significantly lower than in macaques and dogs. Since most of the current experimental mice are SPF mice, whether the *in vivo* experiments using the CRISPR system in mice can simulate humans and other large animals must be concerned. Correspondingly, the antibodies of the baby monkeys are mild, and the early treatment using CRISPR system might have a lower immune response. At present, adaptive immunity against Cas9 has been confirmed in humans, and we have also confirmed the presence of adaptive immunity against Cas protein in monkeys, dogs and mice.

Currently, mammalian gene editing generally serves two purposes. One is to use the CRISPR system to edit the early embryo stage to obtain animal models of certain diseases, and the other is to deliver CRISPR system to specific tissues and organs by vectors such as AAVs to treat certain genetic diseases[8, 9]. Due to ethical issue, direct editing of human embryos is subject to strict ethical review. NHPs are very close to humans because of their genetic homology and physiological functions[31], and are good model to perform experiments that are limited in human.

For gene editing monkeys, the CRISPR system is usually injected at the embryonic stage. The dose of this injection is excessive, and the fate of Cas protein after gene editing is still unknown[4]. Our research shows that no matter what method of gene editing is used, there is no difference in antibody levels between adult gene editing monkeys and the normal population. In mammals, IgG is an immunoglobulin that can cross the placental barrier[40]. Usually, the fetus still has the maternal IgG after birth[41]. Therefore, dynamic monitoring of PAIR-C at different stages is necessary for future *in vivo* gene therapy.

For gene therapy, the CRISPR system is usually packaged into AAV vectors and does not come into direct contact with the antibody, while the primary immune response is the humoral immune response of AAVs and the cellular immune response induced by the Cas proteins. There have been reports of cellular immunity against Cas proteins, but they are all based on *in vitro* cell experiments[19]. After *in vivo* Cas proteins work, it is still unknown whether these cells edited by CRISPR system will activate cellular immunity. For the gene therapy of DMD mice and DMD dogs that have been reported so far, there is no data for immunological monitoring worthy of reference[13–15, 42]. In addition, for gene correction in macaques, if there is a tolerable threshold for Cas proteins still needs to be explore.

In a word, there is still a long way to go for gene editing technology to be applied in clinic. Off-target effect, immune response and toxicity are three major safety problems faced by CRISPR system and other gene editing systems. If these problems are not solved, it will be difficult for gene editing technology to be applied in clinic. The existence of AAVs and Cas proteins antibodies is a problem that must be faced and solved in therapeutic research. There may be a tolerance value between immune rejection and therapeutic effect, which needs to be studied continuously and carefully on NHPs close to human beings. In addition, low immunogenicity and high targeting, high efficiency and low toxicity should be the focus of future research.

## Materials and Methods

### Blood collection and serum separation

Macaques and dogs were collected by venous blood, and SPF mice were bled by eyeballs. Approximately 1 ml blood was drawn into 1.5 ml EP tube, incubated on ice for 2 h, centrifuged at 10,000 g for 10 min. The supernatant was taken to new EP tube and stored at −80 °C. The macaques used in the experiments were approved in advance by the Institutional Animal Care and Use Committee and were performed in accordance with the Association for Assessment and Accreditation of Laboratory Animal Care International for the ethical treatment of primates.

### Construction of prokaryotic expression vector

The gene sequences of SaCas9, SpCas9, AsCas12a, and LbCas12a proteins were derived from pX601-AAV-CMV::NLS-SaCas9-NLS-3xHA-bGHpA;U6::BsaI-sgRNA (Addgene plasmid # 61591), pSpCas9(BB)-2A-GFP (PX458) (Addgene plasmid # 48138), pY010 (pcDNA3.1-hAsCpf1) (Addgene plasmid # 69982), and pY016 (pcDNA3.1-hLbCpf1) (Addgene plasmid # 69988), respectively, which were gifts from Feng Zhang[33–35]. By comparing the fragments of the expression vector and four target genes, we selected SalI and XbaI as the insertion sites between the multiple clone site of the pET28b prokaryotic expression vector. Then primers were designed for the gene sequences of *SaCas9, SpCas9, AsCas12a* and *LbCas12a*. The forward primer 5′ was added with SalI digestion site sequence, and the reverse primer 5′ with XbaI digestion site sequence (S1a Fig). The *SaCas9, SpCas9, AsCas12a* and *LbCas12a* genes were amplified by PCR using pX601, pX458, pY010 and pY016 plasmids as templates. According to the instructions, 10 ng plasmid was used as template, and 30 cycles of reaction were carried out using PrimeSTAR GXL DNA polymerase (Takara, # R050B) (denaturation at 98 °C for 10 seconds, denaturation at 60 °C for 15 seconds, and elongation at 68 °C for 4 minutes). The pET28b plasmid and the PCR amplified target sequences were double digested using SalI-HF (NEB, # R3138S) and XbaI (NEB, # R0145S) restriction endonucleases. The digested product was recovered using EasyPure Quick Gel Extraction Kit (TransGen Biotech, # EG101-02). The vector and the fragment of interest were ligated overnight using T4 DNA ligase (NEB, # M0202T) at 16 °C. The ligation product was transformed into DH5α competent cell (TIANGEN, # CB101), plated on LB solid plates containing kanamycin, and cultured at 37 °C overnight. The transformants were picked, cultured in LB liquid medium containing kanamycin, and shaken overnight at 180 rpm. The next day, use 50% glycerol to keep the bacteria mixed with 1:1 and store at −80 °C.

### Prokaryotic expression

Plasmids of the four constructed expression vectors were extracted and transformed into Rosetta (DE3) competent cell (TIANGEN, # CB108), respectively, and plated on LB solid plates containing kanamycin, then cultured at 37 °C overnight. The transformants were picked and cultured overnight in 5 ml of Lb liquid medium containing kanamycin. The next day, the liquid was inoculated to 100 mL of Lb liquid medium containing kanamycin, cultured at 37 °C, 180 rpm. To OD600 in the range of 0.6-0.8, then add 0.1 mM IPTG, 18 °C, 120 rpm and continue to shake for 20 h. Then, the bacterial fluid was collected by centrifugation at 8000g for 10 minutes, the supernatant was removed, and washed twice with PBS.

### Proteins purification

The protein was purified using HisTrap excel, an immobilized metal affinity chromatography purification column (GE Healthcare, # 17371205), and equilibration buffer (20 mM sodium phosphate, 0.5 M NaCl, pH 7.4), wash buffer (20 mM sodium phosphate, 0.5 M NaCl, 20 mM imidazole, pH 7.4) and elution buffer (20 mM sodium phosphate, 0.5 M NaCl, 500 mM imidazole, pH7.4) were prepared according to the instructions. Filter buffer through a 0.45 μm filter before use. The collected bacteria were sufficiently resuspended with 40 ml equilibration buffer, followed by the addition of 400 μl 100 mM PMSF. The bacteria were lysed using ultrasonic cell disrupter and the lysate was centrifuged at 12,000 g for 10 min. Then the supernatant was filtered using 0.45 μm filter, and the sample was loaded through the prepared HisTrap excel purification column at a flow rate of 1 ml/min. The impurity proteins were washed with 20 column volume (CV) wash buffer at 1 ml/min, followed by 5 CV elution buffer elutes target protein at 1 ml/min.

### Western blot

SaCas9, SpCas9, AsCas12a and LbCas12a proteins were incubated with SDS-PAGE sample loading buffer (Biosharp, # BL502A) at 95 °C for 5 min, and 500 ng protein was loaded to 10% TGX (Tris-Glycine eXtended) Stain-Free FastCast polyacrylamide gels (Bio-rad, # 1610183), 200 V running for 45min. Subsequently, these proteins were transferred to polyvinylidene fluoride (PVDF) membrane (Millipore, # IPVH00010) and blocked with 5% skim milk (BD, # 232100) in TBST for 2 h at room temperature. Blots were incubated with 5% skim milk diluted at 1:200 (Macaque and Dog), 1:100 (SPF mouse) diluted serum at 4 °C overnight. The next day, the blots were washed 5 times with TBST for 3 minutes each time. Subsequently, anti-monkey IgG HRP secondary antibody (Abcam, # ab112767), anti-dog IgG HRP secondary antibody (Bioss, # bs-0303R) and anti-mouse IgG HRP secondary antibody (Millipore, # AP308P) were diluted 1:2000 with 5% skim milk, added to the corresponding blots, respectively, and incubated for 2 h at room temperature on a shaker. Thereafter, the blots were washed 5 times with TBST for 3 minutes each, followed by imaging with Immobilon chemiluminescent HRP substrate (Millipore, # WBKLS0500).

### SpCas9 and AAV-specific antibody test by ELISA

SpCas9 protein was diluted to 2 μg/ml with ELISA coating buffer (Solarbio, # C1055). AAV8 and AAV9 was also diluted to 2×10^10^ vg/ml with ELISA coating buffer. Then 0.1 ml per well was added to 96-well plate (Thermo Fisher Scientific, #3855) respectively, and incubated overnight at 4 °C. The next day, the plate was washed 3 times with PBST, 250 μl 5% skim milk (dissolved in PBST) was added to each well and blocked at 37 °C for 3 hours. Subsequently, it was washed 3 times with PBST, 0.1 ml serum diluted 1:200 for SpCas9 and 1:300 for AAV8/9 were added to the coated reaction well. The mixture was incubated at 37 °C for 1 hour, and then washed with PBST 5 times. Fresh 1:10000 (mouse and dog 1:3000) 5% skim milk diluted enzyme-labeled secondary antibody (anti-Monkey IgG HRP, anti-Dog IgG HRP, anti-Ms IgG HRP) 0.2 ml was added and incubated at 37 °C for 45min, then washed 5 times with PBST. Subsequently, 0.1 ml TMB single-component substrate solution (Solarbio, # PR1200) was added to each well, and the mixture was incubated at 37 °C for 10 minutes. Thereafter, the reaction was terminated by adding 100 ml 1 M sulfuric acid solution to each well. The OD value of each well was measured at 450 nm on microplate reader to indicate the antibody concentration of SpCas9, AAV8 and AAV9 in serum samples.

## Acknowledgments

This work was supported by the Yunnan Key Laboratory of Primate Biomedicine Research. We thank the Laboratory Animal Department’s breeding and management staff for their assistance in this project.

## Supporting information

**S1 Fig.**
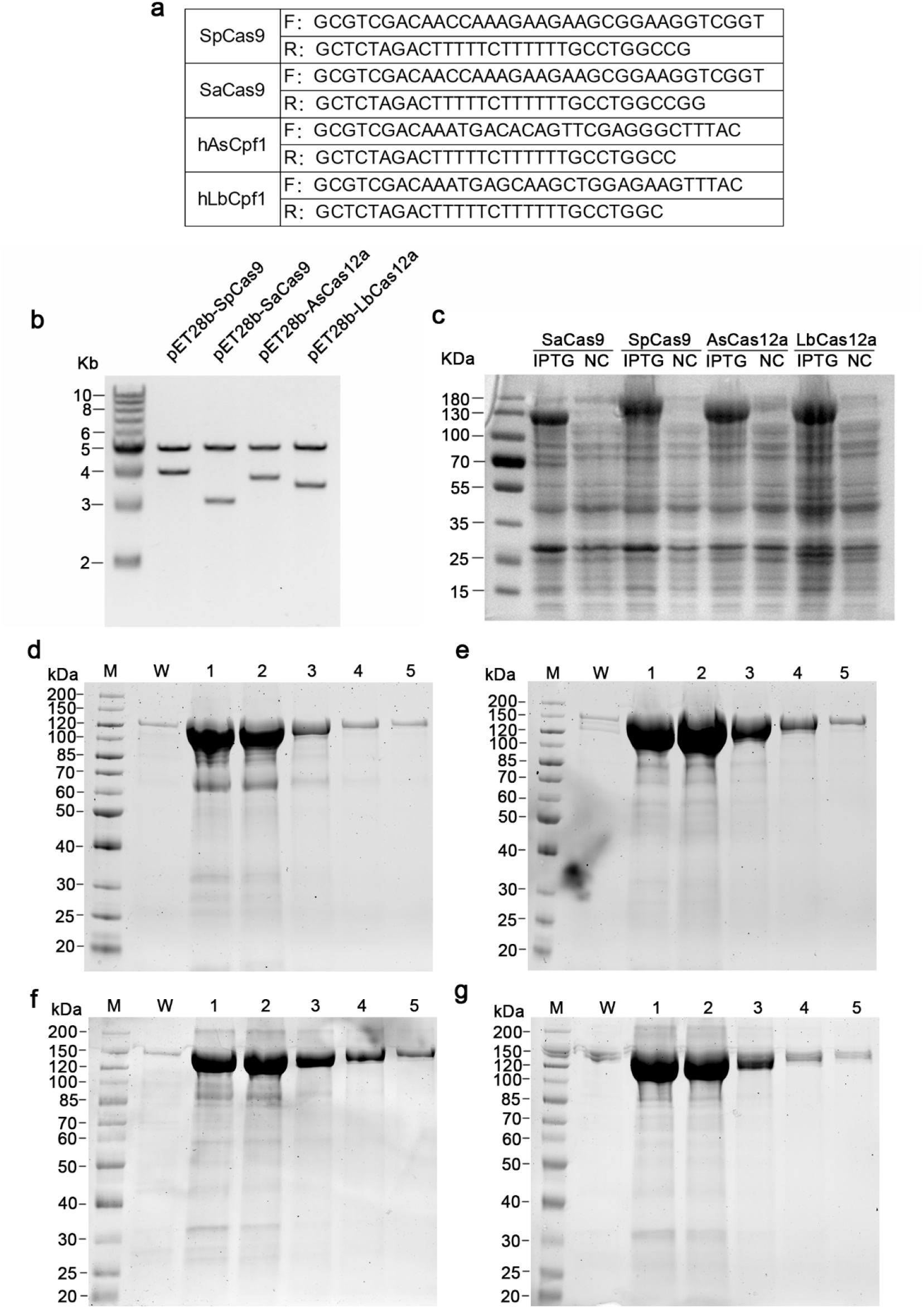
Construction of prokaryotic expression vectors and purification of proteins. (a) Primers used to amplify the sequences of the *SaCas9, SpCas9, AsCas12a* and *LbCas12a* genes. (b) The prokaryotic expression vectors of SpCas9, SaCas9, AsCas12a and LbCas12a proteins were constructed using pET28b, and the successfully constructed expression vectors were verified by double digestion with XbaI and SalI restriction endonucleases. (c) The successfully verified expression vectors were transformed into *E. coli* Rosetta (DE3) strain and the SaCas9, SpCas9, AsCas12a, LbCas12a proteins were expressed under IPTG induction. NC is the result without IPTG induction. (d), (e), (f), (g) are the purification results of SaCas9, SpCas9, AsCas12a and LbCas12a proteins, respectively. M stands for maker, W stands for wash buffer, and 1, 2, 3, 4, 5 are the elution sequences in the elution buffer at a concentration of 500 mM imidazole, respectively.

**S2 Fig.**
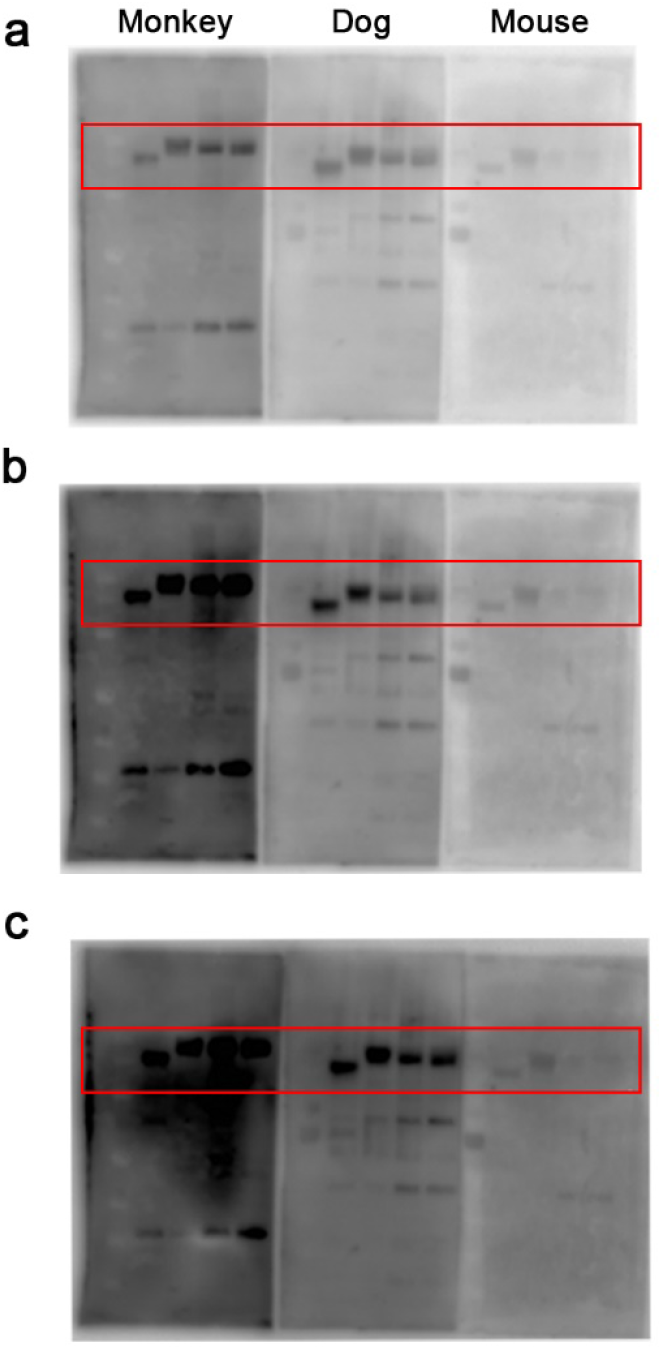
Western blot images of monkey, dog and mouse. (a), (b), and (c) are western blot comparisons of monkey, dog, and mouse exposure for 60s, 180s, and 300s, respectively. From left to right are SaCas9, SpCas9, AsCas12a, LbCas12a.

**S1 Table.**
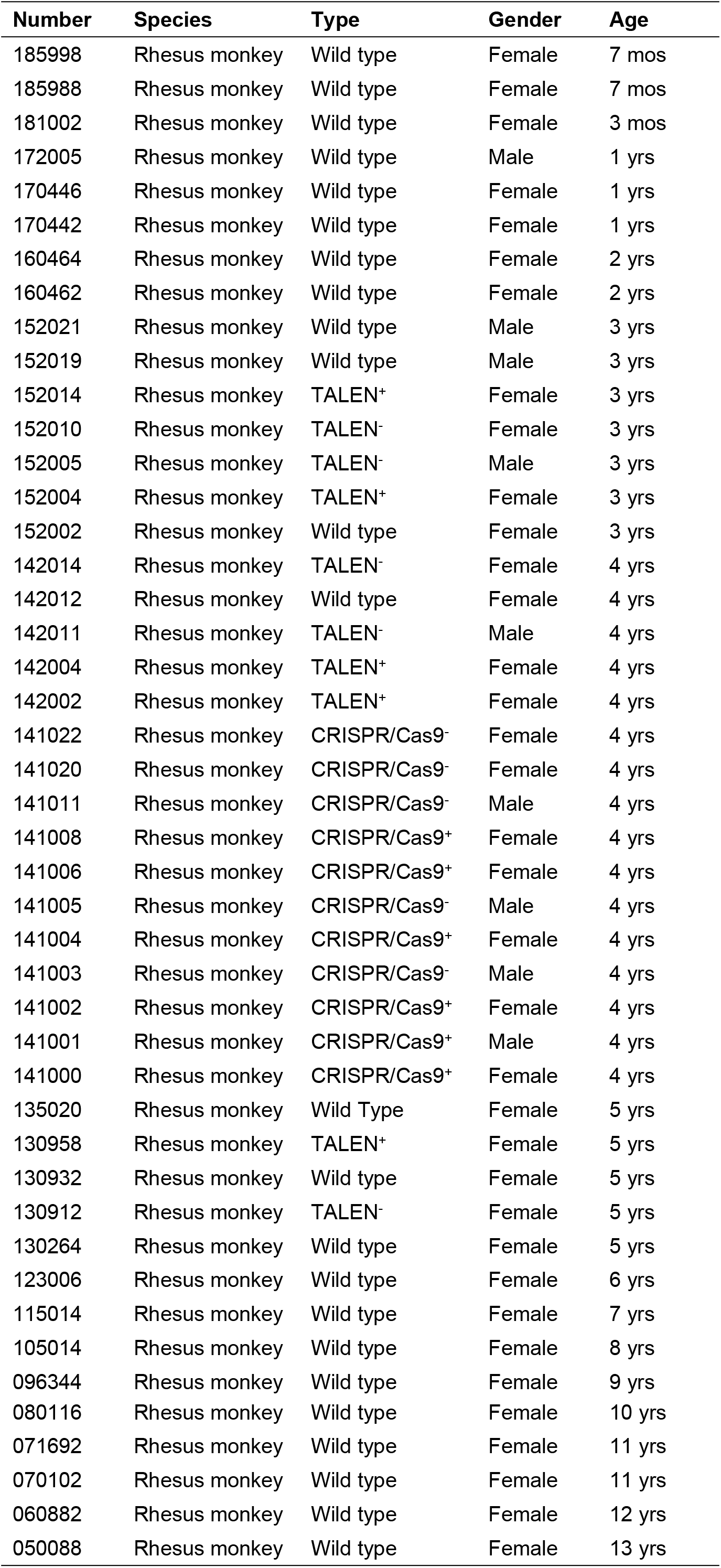
Macaque monkey information statistics for experiments.

| **Number** | **Species** | **Type** | **Gender** | **Age** |
| --- | --- | --- | --- | --- |
| 185998 | Rhesus monkey | Wild type | Female | 7 mos |
| 185988 | Rhesus monkey | Wild type | Female | 7 mos |
| 181002 | Rhesus monkey | Wild type | Female | 3 mos |
| 172005 | Rhesus monkey | Wild type | Male | 1 yrs |
| 170446 | Rhesus monkey | Wild type | Female | 1 yrs |
| 170442 | Rhesus monkey | Wild type | Female | 1 yrs |
| 160464 | Rhesus monkey | Wild type | Female | 2 yrs |
| 160462 | Rhesus monkey | Wild type | Female | 2 yrs |
| 152021 | Rhesus monkey | Wild type | Male | 3 yrs |
| 152019 | Rhesus monkey | Wild type | Male | 3 yrs |
| 152014 | Rhesus monkey | TALEN<sup>+</sup> | Female | 3 yrs |
| 152010 | Rhesus monkey | TALEN<sup>-</sup> | Female | 3 yrs |
| 152005 | Rhesus monkey | TALEN<sup>-</sup> | Male | 3 yrs |
| 152004 | Rhesus monkey | TALEN<sup>+</sup> | Female | 3 yrs |
| 152002 | Rhesus monkey | Wild type | Female | 3 yrs |
| 142014 | Rhesus monkey | TALEN<sup>-</sup> | Female | 4 yrs |
| 142012 | Rhesus monkey | Wild type | Female | 4 yrs |
| 142011 | Rhesus monkey | TALEN<sup>-</sup> | Male | 4 yrs |
| 142004 | Rhesus monkey | TALEN<sup>+</sup> | Female | 4 yrs |
| 142002 | Rhesus monkey | TALEN<sup>+</sup> | Female | 4 yrs |
| 141022 | Rhesus monkey | CRISPR/Cas9<sup>-</sup> | Female | 4 yrs |
| 141020 | Rhesus monkey | CRISPR/Cas9<sup>-</sup> | Female | 4 yrs |
| 141011 | Rhesus monkey | CRISPR/Cas9<sup>-</sup> | Male | 4 yrs |
| 141008 | Rhesus monkey | CRISPR/Cas9<sup>+</sup> | Female | 4 yrs |
| 141006 | Rhesus monkey | CRISPR/Cas9<sup>+</sup> | Female | 4 yrs |
| 141005 | Rhesus monkey | CRISPR/Cas9<sup>-</sup> | Male | 4 yrs |
| 141004 | Rhesus monkey | CRISPR/Cas9<sup>+</sup> | Female | 4 yrs |
| 141003 | Rhesus monkey | CRISPR/Cas9<sup>-</sup> | Male | 4 yrs |
| 141002 | Rhesus monkey | CRISPR/Cas9<sup>+</sup> | Female | 4 yrs |
| 141001 | Rhesus monkey | CRISPR/Cas9<sup>+</sup> | Male | 4 yrs |
| 141000 | Rhesus monkey | CRISPR/Cas9<sup>+</sup> | Female | 4 yrs |
| 135020 | Rhesus monkey | Wild Type | Female | 5 yrs |
| 130958 | Rhesus monkey | TALEN<sup>+</sup> | Female | 5 yrs |
| 130932 | Rhesus monkey | Wild type | Female | 5 yrs |
| 130912 | Rhesus monkey | TALEN<sup>-</sup> | Female | 5 yrs |
| 130264 | Rhesus monkey | Wild type | Female | 5 yrs |
| 123006 | Rhesus monkey | Wild type | Female | 6 yrs |
| 115014 | Rhesus monkey | Wild type | Female | 7 yrs |
| 105014 | Rhesus monkey | Wild type | Female | 8 yrs |
| 096344 | Rhesus monkey | Wild type | Female | 9 yrs |
| 080116 | Rhesus monkey | Wild type | Female | 10 yrs |
| 071692 | Rhesus monkey | Wild type | Female | 11 yrs |
| 070102 | Rhesus monkey | Wild type | Female | 11 yrs |
| 060882 | Rhesus monkey | Wild type | Female | 12 yrs |
| 050088 | Rhesus monkey | Wild type | Female | 13 yrs |

**S2 Table.** Dog information statistics for experiments.

| **Number** | **Species** | **Gender** | **Age** |
| --- | --- | --- | --- |
| D1 | Dog | Male | 1 yrs |
| D2 | Dog | Female | 2 yrs |
| D3 | Dog | Male | 1 yrs |
| D4 | Dog | Female | 4 yrs |
| D5 | Dog | Male | 1 yrs |
| D6 | Dog | Female | 2 mos |
| D7 | Dog | Female | 1 yrs |
| D8 | Dog | Male | 2 yrs |
| D9 | Dog | Female | 8 yrs |
| D10 | Dog | Male | 4 mos |

**S3 Table.**
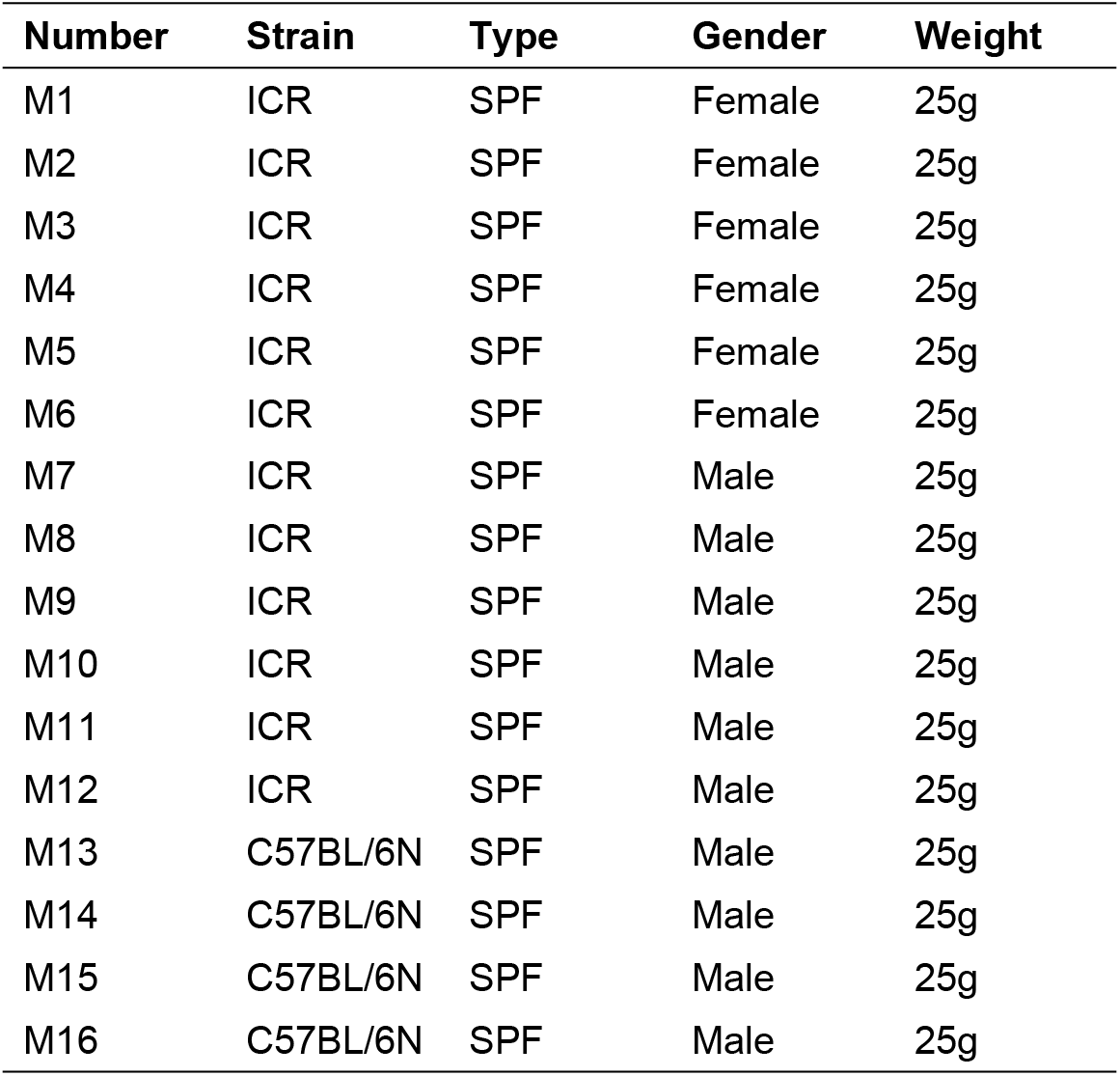
Mouse information statistics for experiments.

| **Number** | **Strain** | **Type** | **Gender** | **Weight** |
| --- | --- | --- | --- | --- |
| M1 | ICR | SPF | Female | 25g |
| M2 | ICR | SPF | Female | 25g |
| M3 | ICR | SPF | Female | 25g |
| M4 | ICR | SPF | Female | 25g |
| M5 | ICR | SPF | Female | 25g |
| M6 | ICR | SPF | Female | 25g |
| M7 | ICR | SPF | Male | 25g |
| M8 | ICR | SPF | Male | 25g |
| M9 | ICR | SPF | Male | 25g |
| M10 | ICR | SPF | Male | 25g |
| M11 | ICR | SPF | Male | 25g |
| M12 | ICR | SPF | Male | 25g |
| M13 | C57BL/6N | SPF | Male | 25g |
| M14 | C57BL/6N | SPF | Male | 25g |
| M15 | C57BL/6N | SPF | Male | 25g |
| M16 | C57BL/6N | SPF | Male | 25g |

**S4 Table.** Statistics of western blot results in macaques.

| **Number** | **SaCas9** | **SpCas9** | **AsCas12a** | **LbCas12a** |
| --- | --- | --- | --- | --- |
| 185998 | + | - | + | + |
| 185988 | + | - | - | + |
| 181002 | + | + | - | - |
| 172005 | + | + | + | + |
| 170446 | + | + | + | + |
| 170442 | + | + | + | + |
| 160464 | + | + | + | + |
| 160462 | + | + | + | + |
| 152021 | + | + | + | + |
| 152019 | + | + | + | + |
| 152014 | + | + | + | + |
| 152010 | + | + | + | + |
| 152005 | + | + | + | + |
| 152004 | + | + | + | + |
| 152002 | + | + | + | + |
| 142014 | + | + | + | + |
| 142012 | + | + | + | + |
| 142011 | + | + | + | + |
| 142004 | + | + | + | + |
| 142002 | + | + | + | + |
| 141022 | + | + | + | + |
| 141020 | + | + | + | + |
| 141011 | + | + | + | + |
| 141008 | + | + | + | + |
| 141006 | + | + | + | + |
| 141005 | + | + | + | + |
| 141004 | + | + | + | + |
| 141003 | + | + | + | + |
| 141002 | + | + | + | + |
| 141001 | + | + | + | + |
| 141000 | + | + | + | + |
| 135020 | + | + | + | + |
| 130958 | + | + | + | + |
| 130932 | + | + | + | + |
| 130912 | + | + | + | + |
| 130264 | + | + | + | + |
| 123006 | + | + | + | + |
| 115014 | + | + | + | + |
| 105014 | + | + | + | + |
| 096344 | - | + | + | + |
| 080116 | + | + | + | + |
| 071692 | + | + | + | + |
| 070102 | + | + | + | + |
| 060882 | + | + | + | + |
| 050088 | + | + | + | + |

**S5 Table.** Statistics of western blot results in dogs.

| **Number** | **SaCas9** | **SpCas9** | **AsCas12a** | **LbCas12a** |
| --- | --- | --- | --- | --- |
| D1 | + | + | + | + |
| D2 | + | + | + | + |
| D3 | + | + | + | + |
| D4 | + | + | + | + |
| D5 | + | + | + | + |
| D6 | + | + | + | + |
| D7 | + | + | + | + |
| D8 | + | + | + | + |
| D9 | + | + | + | + |
| D10 | + | + | + | + |

**S6 Table.** Statistics of western blot results in mice.

| **Number** | **SaCas9** | **SpCas9** | **AsCas12a** | **LbCas12a** |
| --- | --- | --- | --- | --- |
| M1 | + | + | + | + |
| M2 | + | + | + | + |
| M3 | + | + | + | + |
| M4 | + | + | + | + |
| M5 | + | + | + | + |
| M6 | + | + | + | + |
| M7 | + | + | + | + |
| M8 | + | + | + | + |
| M9 | + | + | + | + |
| M10 | + | + | + | + |
| M11 | + | + | + | + |
| M12 | + | + | + | + |
| M13 | + | + | + | + |
| M14 | - | - | - | - |
| M15 | + | + | + | + |
| M16 | + | + | + | + |

**S1 Data. Western blot images of serum antibodies against four Cas proteins in macaques, dogs and mice.**

